# Temporally-precise disruption of prefrontal cortex informed by the timing of beta bursts impairs human action-stopping

**DOI:** 10.1101/843557

**Authors:** Ricci Hannah, Vignesh Muralidharan, Kelsey K Sundby, Adam R Aron

## Abstract

Human action-stopping is thought to rely on a prefronto-basal ganglia-thalamocortical network, with right inferior frontal cortex (rIFC) posited to play a critical role in the early stage of implementation. Here we sought causal evidence for this idea in experiments involving healthy human participants. We first show that action-stopping is preceded by bursts of electroencephalographic activity in the beta band over prefrontal electrodes, putatively rIFC, and that the timing of these bursts correlates with the latency of stopping at a single-trial level: earlier bursts are associated with faster stopping. From this we reasoned that the integrity of rIFC at the time of beta bursts might be critical to successful stopping. We then used fMRI-guided transcranial magnetic stimulation (TMS) to disrupt rIFC at the approximate time of beta bursting. Stimulation prolonged stopping latencies and, moreover, the prolongation was most pronounced in individuals for whom the pulse appeared closer to the presumed time of beta bursting. These results help validate a model of the neural architecture and temporal dynamics of action-stopping. They also highlight the usefulness of prefrontal beta bursts to index an apparently important sub-process of stopping, the timing of which might help explain within- and between-individual variation in impulse control.

## INTRODUCTION

Studies using causal methods, such as lesions in human patients and brain stimulation, have shown that the right inferior frontal cortex (rIFC) in humans is a critical node in a putative prefronto-basal ganglia-thalamocortical network for stopping actions (Aron et al. 2003; Chambers et al. 2006, 2007; Verbruggen et al. 2010; Kohl et al. 2019). The rIFC is thought to play a role early in the implementation of the stop process, potentially initiating it via the hyper-direct pathway to the subthalamic nucleus (Aron et al. 2014). Electrophysiological studies of action-stopping using the Stop Signal Task (Verbruggen and Logan 2009) – which requires the cancellation of a prepotent action when a “stop signal” occurs – seem consistent with this early role. For example, electrocorticographic activity from rIFC shows an increase in beta band oscillatory power in successful versus failed stop trials, and moreover in the time-period between the stop signal and the end of the inferred stop process (Swann et al. 2009; Wessel, Conner, et al. 2013). A similar pattern of beta band power is also seen in scalp electroencephalography (EEG) and magnetoencephalography (MEG) studies, putatively related to the same rIFC ‘generator’ (Wagner et al. 2018; Castiglione et al. 2019; Schaum et al. 2020). However, there is a lack of causal evidence showing *when in time* activation of rIFC is critical for stopping. Knowing precisely when rIFC is involved in stopping is fundamental to establishing the temporal dynamics of the wider stopping network (Allen et al. 2018; Schaum et al. 2020), and for more general questions of prefrontal-basal ganglia interactions and cognitive control that we will return to below.

Here we delivered single pulse transcranial magnetic stimulation (TMS) over rIFC during a stop signal task. We aimed to a) transiently disrupt rIFC with precisely-timed pulses and b) assess the behavioral impact of that disruption with a recently-validated single-trial readout of stopping latency.

We based our timing for stimulation on recent findings about beta oscillations from putative rIFC. Whereas the above-cited studies refer to average beta band power, we have also recently shown how this is underpinned by transient beta bursts (Jana et al. 2019). Here, we reasoned that the integrity of rIFC at the time of these bursts is critical to the successful implementation of the stop process, and that disrupting it at this time would therefore impair stopping.

To evaluate the behavioral impact of rIFC disruption we used our recently-validated measure of stopping latency derived from electromyographic (EMG) recordings during the Stop Signal Task (Jana et al. 2019). The measure relies on the fact that even trials in which participants successfully stop a button-press sometimes contain small bursts of EMG presumably reflecting voluntary commands to move. Importantly, however, these EMG bursts are cut short ∼160 ms after the stop signal (Raud and Huster 2017; Jana et al. 2019), and we proposed that this reflected the timing of the implementation of the stop process at the muscle. The idea was supported by evidence that the time of EMG cancellation (CancelTime) coincides temporally with a widespread, motor system suppression (Jana et al., 2019; see also Coxon et al., 2006; Badry et al., 2009; Wessel et al., 2013), perhaps mediated via the subthalamic nucleus (Wessel et al. 2016), which suggests an active withdrawal/inhibition of motor drive. Here we use this CancelTime metric as a single trial estimate of stopping latency.

We report results from several experiments. For Experiment 1 we re-analyzed existing EEG data to test whether the time of right frontal beta bursts (BurstTime) relate to CancelTime at the single-trial level. For Experiment 2 we ran fMRI to localize the rIFC in each participant, and also analyzed EEG from a second session in these participants to confirm the timing of beta bursts in this new population. Then, we studied these same participants in Experiment 3, involving event-related fMRI-guided TMS. We used the information about the timing of the bursts as a proxy for stop-related rIFC activity in this sham-controlled and double-blind TMS experiment.

## MATERIALS AND METHODS

We pre-registered our methodological plan for Experiment 3, specifically the number of participants, our approach to double-blinding, and our specific hypotheses and statistical analyses (see URL #1 at end of manuscript).

### Participants

Participants were healthy, human volunteers who provided written informed consent and were compensated $20/hour for their time. The experiments were approved by the UCSD Institutional Review Board.

#### Experiment 1 (EEG)

Fifteen participants (9 females; age 21±0.4 years; all right-handed) were recruited as part of a separate study (Jana et al. 2019). Here we present data based on a new set of analyses. Note that two participants were excluded: one due to misaligned EEG markers resulting from a technical issue and another because we could not identify a right frontal spatial filter based on our standard method (Wagner et al. 2018; Castiglione et al. 2019).

#### Experiment 2 (EEG and functional magnetic resonance imaging)

Thirty-six participants were recruited as part of a separate study involving, in separate sessions, functional magnetic resonance imaging (fMRI) and EEG recordings during the Stop Signal Task. Of these, 21 subsequently returned to participate in the TMS experiment (Experiment 3; see below), which used the fMRI scans to localize stimulation to the rIFC. One participant was excluded (see next section). Only data from the final 20 participants included in Experiment 3 are presented here. EMG data were also unavailable for one participant as this element of the project was added after they had participated, and another because a reliable right frontal EEG component could not be determined, so n = 19 for EMG measurements and n = 18 for the EEG-EMG aspect of the experiment.

#### Experiment 3 (TMS)

21 participants (11 females; age 19 ± 0.4 years; 19 right-handed, 2 left-handed) with no contraindications to TMS (Rossi et al. 2011) were recruited, but one was excluded because we were unable to adequately co-register the brain scan in the TMS neuronavigation system.

### Stop Signal Task

The task was coded in MATLAB 2016b (Mathworks, USA) and presented using Psychtoolbox (Brainard 1997). Trials began with a fixation (white square) being presented at the center of the screen for 500 ± 50 ms (Figure 1a). This was then replaced by an imperative cue (white arrow), indicating which digit participants were to respond with: a left facing arrow required them to press a key by abducting their right index finger, and a right facing arrow required them to press a key by flexing their little finger. Participants were encouraged to respond as fast and accurately as possible. The stimuli remained on the screen for 1 s. If participants did not respond within this time, the trial aborted, and ‘Too Slow’ was presented. Trials like these, that simply required a response, are referred to as “Go” trials and accounted for 75% of all trials. In the other 25% of trials, the white arrow turned red after a variable stop signal delay (SSD), and participants were required to try and prevent a response (Stop trials). The SSD was adjusted depending on the success or failure of stopping a response using two independent staircases (for right and left directions), where the SSD increased and decreased by 50 ms following a Successful Stop and Failed Stop, respectively. Each trial was followed by an inter trial interval (ITI) and the entire duration of each trial including the ITI was 2.5 - 4.5 s depending on the experiment.

**Figure 1.**
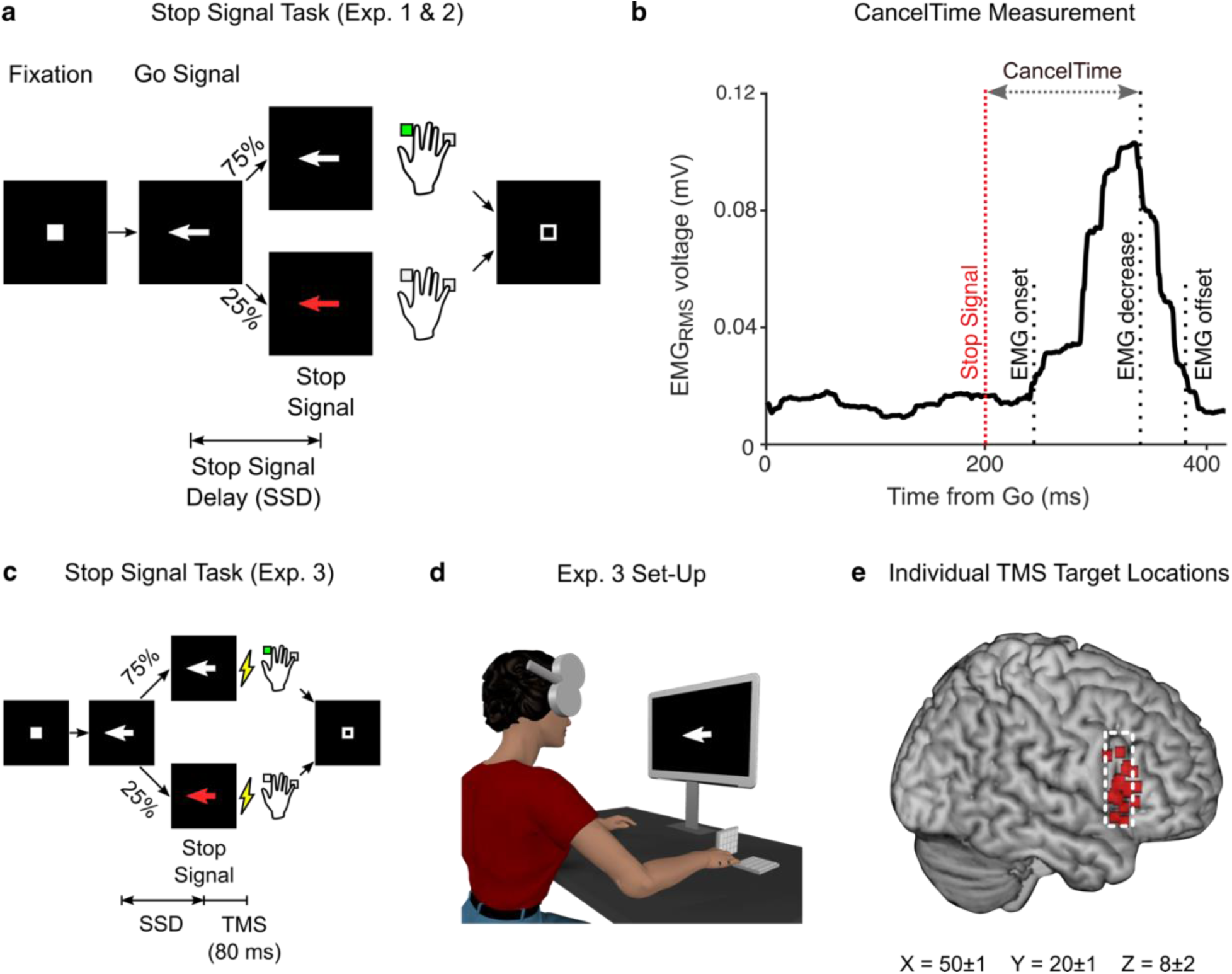
Methods for all Experiments. (a) Trial structure for the Stop Signal Task in experiments 1 and 2. (b) Example EMG recording from a single muscle depicting an EMG burst on a Successful Stop trial where no overt key press was recorded. The time between the stop signal and EMG decrease reflects CancelTime. (c) Trial structure for the Stop Signal Task in experiment 3, depicting the time of TMS stimuli 80 ms after the stop signal. (d) Set-up for the TMS experiment showing TMS location over the rIFC target, with key pads for recording key press responses. (e) Individual TMS target locations projected onto the brain surface over the pars opercularis region (dashed white line) of the inferior frontal gyrus, based on fMRI. Mean coordinates are in Montreal Neurological Institute space.

### Electromyography (EMG)

Surface EMG recordings were made from the right first dorsal interosseous (FDI) and abductor digiti minimi (ADM) muscles. EMG signals were amplified ×5000 between 30-1000 Hz (Grass QP511 AC amplifier, Grass Instruments, West Warwick, USA), digitized at 1000 Hz (Micro 1401 mk II, Cambridge Electronic Design, Cambridge, UK) and recorded via data acquisition software (Signal version 4, Cambridge Electronic Design, Cambridge, UK).

### Electroencephalography (EEG)

In both Experiments 1 and 2, 64 channel EEG was recorded in the standard 10/20 configuration (Easycap and BrainVision actiCHamp amplifier, Brain Products Gmbh, Gilching, Germany) and digitized at 1000 Hz.

### Functional magnetic resonance imaging (MRI)

In order to ensure accurate and reproducible positioning of the TMS coil over the rIFC target in Experiment 3, we used participants’ fMRI scans from Experiment 2 combined with neuronavigation software (Brainsight, Rogue Research Inc., Montreal, Canada). Participants were scanned using a 3T GE scanner at the Center for Functional Magnetic Resonance Imaging at the University of California San Diego. Each scanning session included an anatomical T1 scan and two 6 minute blocks of the stop signal task that reliably activates our target region, the pars opercularis of the rIFC (Aron and Poldrack 2006). 182 functional T2-weighted echoplanar images were acquired for each of the task blocks (TR, 2 s). Data were preprocessed using Functional Magnetic Resonance Imaging of the Brain (FMRIB) Software Library (FSL) (www.fmrib.ox.ac.uk/fsl). Analysis methods were exactly as in Aron and Poldrack (2006), except that the final activation map for each participant contrasted all stop trials (Successful and Failed) versus Go trials (instead of Successful Stop trials vs Go trials), in order to have more observations. The resulting activation map was back-registered onto each participant’s native space, and then masked with the pars opercularis from the Harvard Oxford Atlas (https://fsl.fmrib.ox.ac.uk/fsl/fslwiki/Atlases). The masked activation was then imported into the neuronavigation software. We selected our stimulation site by identifying the site within the pars opercularis mask that exhibited peak activation related to stopping. We held the TMS coil so that the TMS-induced currents were perpendicular to the orientation of the pre-central sulcus. During the experiment, TMS coil position and orientation relative to the target site were recorded in real time for each pulse delivered.

### Transcranial magnetic stimulation (TMS)

For Experiment 3, single pulses were delivered over the right pars opercularis target via a TMS device (PowerMag Lab 100, MAG&More GMBH, Munich, Germany) delivering full sine wave pulses, and connected to one of two 70 mm figure-of-eight coils: real coil (Double coil PMD70-pCool) or sham coil (Double coil PMD70-pCool-Sham). The sham coil is identical in appearance aside from labelling at the distal end of the coil. This labelling was covered during the experiment to ensure the experimenter remained blind to the coil being used. The coil elicits sounds and scalp sensations (e.g. via superficial stimulation which activates scalp muscles) that are very similar to those elicited by real stimulation, thus providing for an effective sham.

Prior to the experiment, the motor hot spot for the right FDI muscle was determined as the position on the scalp where slightly supra-threshold stimuli produced the largest and most consistent MEPs. The coil was positioned on the scalp over the left primary motor cortex representation of the FDI muscle and oriented so that the coil handle was approximately perpendicular to the central sulcus, *i.e.* at ∼45° to the mid-sagittal line, and the initial phase of current induced in the brain was posterior-to-anterior across the central sulcus. Resting motor threshold (RMT) was defined as the lowest intensity to evoke an MEP of at least 0.05 mV in 5 of 10 consecutive trials while participants were at rest. We then established the stimulus intensity to be used during task for the TMS_Real_ coil, which was set to 120% RMT. The real TMS coil was them moved to the pars opercularis target site (Figure 1c and 1d) and participants received three stimuli at 120% RMT, separated by 5 seconds. They were asked at the end to rate the overall intensity of the sensations elicited by the stimuli on a scale of 0-10. Following this, the TMS_Sham_ coil was attached and positioned over the same site. Participants received three stimuli in the same manner, and at the same stimulator output, and were asked to rate the intensity of sensations. If the rating was lower or higher than that for TMS_Real_ the stimulator output was adjusted up or down, respectively, by 1% of maximum stimulator output until the perceptual ratings matched that of TMS_Real_, and this output was used for TMS_Sham_ in the main experiment. Using this approach, resting motor threshold was 48 ± 2 % stimulator output and thus the intensity of TMS_Real_ during the main experiment was 58 ± 2 % stimulator output. With this stimulator output, the mean perceptual rating of stimulation intensity was 6.4 ± 0.3 out of 10. In order to match the perceptual ratings between TMS_Real_ and TMS_Sham_, it was often necessary to increase the stimulus intensity of TMS_Sham_ (62 ± 2 % stimulator output) such that it was slightly higher compared to TMS_Real_.

Participants were not explicitly told that two coils were being used, nor that one of them was a sham coil. In order to ensure the experimenter remained blind to the TMS coil being used, the stimulator output and the labelling on the coil were both covered up and the coils were relabeled A and B, randomly, by another researcher not directly involved in the study. This person set the intensity and changed coils throughout the experiment. Participants and the experimenter also wore ear plugs during the experiment to mask the noise elicited by each coil. Participants completed a written debrief at the end of the experiment asking: (1) if they noticed any changes in the sensations elicited by TMS throughout the experiment; and (2) if so, to describe the intensity, location and quality of those changes. The experimenter also documented which coil they thought was real and which sham.

### Experimental Design

Following an initial block of practice trials, participants in each experiment completed the Stop Signal Task as follows:

#### Experiment 1

24 blocks of 80 trials each (1440 Go trials and 480 Stop trials) with concurrent EEG recording. Here we tested the hypothesis that BurstTime and CancelTime, as our measures of the timing of right frontal cortical activity and stopping latency, would correlate with one another on a trial-by-trial basis.

#### Experiment 2

Participants completed the task on two separate days. On day one, the task was performed in the MRI scanner and participants completed 2 blocks of 256 trials (384 Go and Stop 128 trials). On day two, the task was performed with concurrent EEG recording and participants completed 4 blocks of 80 trials (240 Go trials and 80 Stop trials). The data here provided us with fMRI for TMS localization in Experiment 3, and also an estimate of BurstTime in the same participants which helped guide the timing of TMS.

#### Experiment 3

12 blocks (6 TMS_Real_ and 6 TMS_Sham_), with each block consisting of 96 trials each (432 Go trials and 144 Stop trials in total for each of TMS_Real_ and TMS_Sham_ conditions). Blocks were grouped in threes, so that the experiment started with either three blocks of TMS_Real_ or TMS_Sham_, decided randomly for each participant. After that, blocks proceeded in groups of three, alternating between TMS_Real_ and TMS_Sham_. A single TMS pulse was delivered in 50% go trials and 100% of stop trials. Given that BurstTime reflects the time of the peak amplitude of the beta burst, and the effects of a single TMS pulse on cortical function and behavior last anywhere from tens of milliseconds up to several hundred (Amassian et al. 1989; Day et al. 1989; Inghilleri et al. 1993; Säisänen et al. 2008; Cash et al. 2017), we estimated that the optimal time to deliver a TMS pulse should be slightly earlier than BurstTime (∼125 ms) such that it caught the rise of the burst (burst duration is ∼200 ms (Jana et al. 2019). We therefore decided to deliver TMS 80 ms after the stop signal (Figure 1e). In go trials, TMS pulses were delivered at the same time as in the most recent stop trial.

Our pre-registered predictions (see URL #1) in brief, were: (a) TMS_Real_ would delay stopping relative to TMS_Sham_, and that this would be evident as an increase in CancelTime and the standard measure of stopping, SSRT; (b) TMS_Real_ would not affect response times (i.e. key press response times or EMG onsets) on Go trials relative to TMS_Sham_, or that because of a potential “braking” function of rIFC (Wessel, Conner, et al. 2013) it would speed up response times; and (c) the effect of TMS_Real_ on both response times and stopping times would be dependent on the time at which the pulse was given on a particular trial relative to the event (i.e. response/stop onset) itself.

### Data analysis

All analyses were performed using MATLAB (R2016b). Note too that for Experiment 3 all data and statistical analyses (see below) were performed blind to TMS coil used in each block, with unblinding only occurring after the completion of statistical analyses. They were also performed in accordance with our pre-registered methods (see URL #1), except in relation to the hypothesis (c) noted above (see EMG-based measurement predictions1b and 2a/b in URL #1). In attempting the latter, we realized there was a logical problem in evaluating the effect of a TMS pulse on response times as a function of the time at which TMS was delivered relative to the response onset, i.e. we can’t know what the response time on that trial would have been had we not delivered the pulse. We therefore shelved this within-participant approach and instead adopted a between-participant approach, hypothesizing that participants with a shorter stopping latency would receive TMS closer to the implementation of the stop and therefore experience a greater disruption than those with a longer stopping latency.

#### Stop Signal Task behavior

Reaction times were determined for both Go and Failed Stop trials. SSRT was computed with the integration method (Verbruggen et al. 2019).

#### EMG data

EMG data were analyzed as described in our recent paper (Jana et al. 2019), the code for which is openly accessible (see URL #2). Briefly, EMG data were filtered using 4_th_ order Butterworth filter to remove 60 Hz line noise. The root-mean square (RMS) of the signal was computed using a centered window of 50 ms. Peaks of EMG activity were identified on a trial-by-trial basis as activity exceeding the mean baseline EMG (determined in the period between the fixation and Go signal) by more than 8 standard deviations. Starting from that peak, we then backtracked to identify the onset of EMG activity as the point where it dropped below 20% of the peak for 5 consecutive ms, and the time between the Go signal this point denoted EMG onset time (Figure 1b). The time when EMG started to decrease was determined as the time following the peak where activity decreased for 5 consecutive ms. CancelTime was measured in Successful Stop trials as the time interval between the stop signal and the decline in EMG activity (Figure 1b). Peak EMG amplitudes often differed for the FDI and ADM muscles, therefore EMG activity was normalized to the average EMG amplitude on Go trials for each muscle separately.

#### EEG analyses

We used EEGLAB (Delorme and Makeig 2004) and custom-made scripts to analyze the data. The data were down-sampled to 512 Hz and band-pass filtered between 2-100 Hz. FIR notch filters were applied to remove line noise (60 Hz) and its harmonics (120 and 180 Hz). EEG data were then re-referenced to the average. The continuous data were visually inspected to remove bad channels and noisy stretches. We then performed logistic Infomax ICA on the noise-rejected data to extract independent components (ICs) for each participant separately (Bell and Sejnowski 1995). Using the DIPFIT toolbox in EEGLAB (Oostenveld and Oostendorp 2002; Delorme and Makeig 2004), we then computed the best-fitting single equivalent dipole matched to the scalp projection for each IC. ICs corresponding to eye movements, muscle, and other non-brain sources were then subtracted from the data. These non-brain ICs were identified using the frequency spectrum (increased power at high frequencies), scalp maps (activity outside the brain) and the residual variance of the dipole (greater than 15%).

#### Selecting a right frontal spatial filter

For both Experiments 1 and 2 we aimed to analyze right frontal beta bursts and their relationship to stopping. To ‘localize’ these bursts and to increase the signal to noise ratio for analysis, we identified a right frontal spatial filter (i.e. a weighting over the 64 electrodes) for each participant.

In Experiment 1, we identified a right frontal spatial filter or, in a few cases where a right frontal filter was not evident, a frontal spatial filter, using a similar Independent Components Analysis approach as in our earlier reports (Wagner et al. 2018; Castiglione et al. 2019). The spatial filter was first identified using the scalp maps. The channel data were then projected onto the identified filter and epoched from −1500ms to 1500ms with respect to the stop signal, only for Successful Stop trials. The identified IC was then validated by confirming an increase in beta power (13-25 Hz) averaged across trials between stop signal and SSRT (with respect to a baseline prior to the Go cue) in the time-frequency map of Successful Stop trials. Therefore, the metric used to validate the IC was independent of our metric of interest which was the time of individual beta bursts within a trial. The time-frequency maps were estimated using Morlet wavelets from 4 to 30 Hz with 3 cycles at low frequencies linearly increasing by 0.5 at higher frequencies. The data were then further epoched into Failed Stop and Go trials for a similar time interval (−1500ms to 1500ms with respect to the Stop signal and Go signal, respectively). Finally, we also identified the peak beta frequency for a participant from the time-frequency map of the Successful stop trials, i.e. the beta frequency with the highest power between stop signal and SSRT. This was then utilized for the beta bursts computation.

In Experiment 2, we derived the spatial filter using a guided source separation approach called generalized eigen decomposition (GED) (for earlier uses of the approach see Parra and Sajda, 2004; Cohen, 2017; Muralidharan et al., 2019) because the number of data points were too low (12 mins of Stop Signal task) to perform the ICA. GED derives a spatial filter which maximizes the SNR of one condition compared to another. In this case GED took advantage of two time periods within Successful Stop trials. Specifically, we contrasted stopping-related activity (i.e. occurring after the stop signal until SSRT) with activity before the Go cue (baseline). To do this we first epoched the data from −2000 ms to 500 ms with respect to the stop signal for Successful stop trials. We then filtered the data around beta frequency (from 10-25 Hz, since right frontal spatial maps were sometimes observed at slightly lower frequencies than beta) using a frequency domain Gaussian kernel having a full width half maximum of 5 Hz. Two time windows were then selected, a stop window (0 to 250 ms) and a baseline window (−1500 to −1000 ms with respect to the Stop signal). We then performed GED for the stop window versus the baseline window, for each frequency separately, deriving a set of spatial filters. A participant-specific spatial filter needed to meet several criteria. First, scalp topography: we selected the component if the scalp map was right frontal (if not right then a frontal topography). Second, it had to occur in the top 6 components, where the components obtained for each frequency were ranked in the order of decreasing variance. Once we identified candidate filter/s using the above criteria, similar to our approach in Experiment 1, the final validation of the filter was based on the time-frequency map of Successful Stop trials. The channel data were projected onto the candidate filter/s and epoched from −1500ms to 1500ms with respect to the stop signal for Successful Stop trials in order to confirm an increase in beta power. When more than one filter was identified then the one with the highest beta power increase between stop signal and SSRT was selected for further analysis.

Having selected a spatial filter for each participant, the data were then epoched into Failed Stop and Go trials for the same time interval as Successful Stops (−1500ms to 1500ms with respect to the Stop signal and Go signal, respectively). As before, in each participant, the beta frequency which had the maximum power between stop signal and SSRT was used for the beta bursts computation.

#### Extracting Beta Bursts

Beta bursts were identified from the epoched data, by filtering the data first at the estimated peak beta frequency (frequency-domain Gaussian, fwhm = 5Hz) for that participant and then defining a burst threshold using the beta amplitude from a baseline period (−1000ms to −500ms with respect to the stop signal in Stop trial; mean SSD – 1000ms to mean SSD – 500ms in Go trials). The complex analytic time series was then obtained using a Hilbert transform. We computed the beta amplitude by taking the absolute of the analytic signal. Subsequently, we quantified the median and standard deviation (SD) of the baseline beta amplitude pooled across all trial types and used it to define the burst threshold. A burst was any period of increase in beta amplitude within a trial that exceeded median + 1.5SD. For each detected burst, the time of the peak beta amplitude was marked as the BurstTime and the burst width/duration was computed using a slightly lower threshold (median + 1 SD) than that used to identify a burst. We marked all the times where the beta amplitude crossed this lower threshold in order to compute burst probability (burst %) across trials. Mean BurstTime for a given period of interest (before and after stop) was calculated for each participant using all trials containing bursts in that period of time, whereas for trial-by-trial correlations of BurstTime with CancelTime we considered only those trials where there was both a beta burst and an EMG burst.

### Statistical Analyses

The majority of data were evaluated with paired *t*-tests and the Bonferroni Correction was applied to control the family-wise error rate when multiple comparisons were performed. Effect sizes were interpreted as small (Cohen’s *d*: 0.2-0.5; Bayes Factor in favor of the alternate hypothesis, *BF_10_*: 1-3), medium (*d*: 0.5-0.8; *BF_10_*: 3-10) or large (*d* > 0.8; *BF_10_* > 10). For comparisons across multiple levels, repeated-measures ANOVA was used. Effect sizes for ANOVAs were interpreted as small (partial eta-squared, *η_p2_*: 0.01-0.06), medium (*η_p2_*: 0.06-0.14), and large (*η_p2_* > 0.14). For correlational analyses, Pearson’s correlation coefficient (*r*) was used. For the trial-by trial correlational analysis between BurstTime and CancelTime, we used the repeated measures correlation method to estimate the common linear association across multiple within-participant correlations (Bakdash and Marusich 2017). All data are presented as mean± SEM.

## RESULTS

### Experiment 1 (EEG experiment): Trial-by-trial beta bursts relate to CancelTime

In this experiment we revisited EEG data from an earlier study (reported in Jana et al., 2019) and analyzed them in a new way to test the relationship between right frontal beta bursts and the single trial metric of action-stopping (so called CancelTime). As reported in the previous study, behavioral performance on the Stop Signal Task was typical (Table 1). The key findings from the previous study, were that the burst probability was higher in Successful Stop trials compared to baseline and to Go trials, and that the average time of the bursts (BurstTime) preceded the time of the EMG decrease (CancelTime). Finally, BurstTime was positively correlated with CancelTime across participants. These data were consistent with the idea that right frontal cortex activity precedes the end of the stop process, a necessary condition for such activity to causally contribute to stopping. This motivated our current attempt to analyze the within-participant, trial-by-trial relationship between BurstTime and CancelTime, which would provide convincing evidence for the notion that the prefrontal beta bursts index the onset of a stop-related process.

**TABLE 1:**
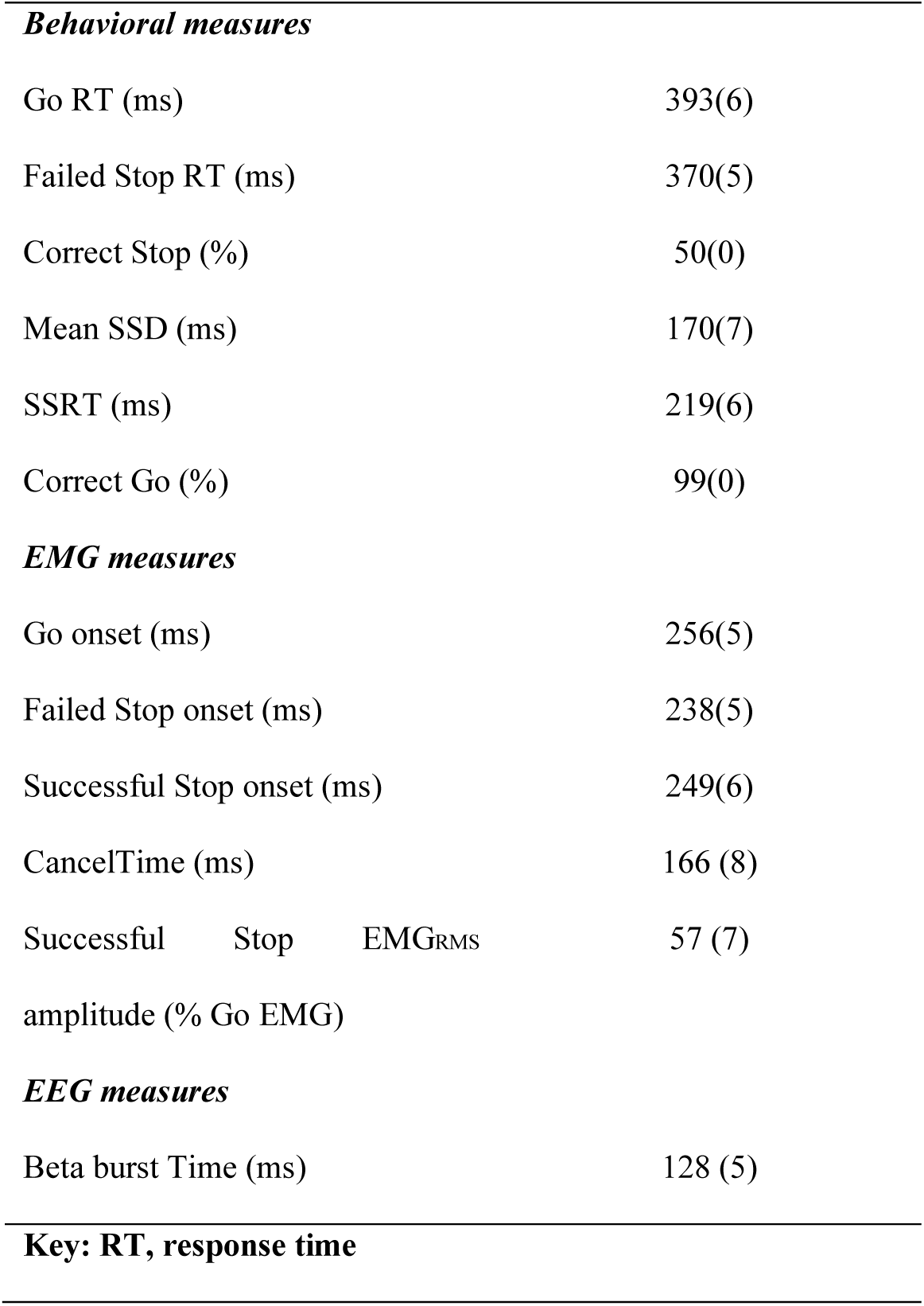
Behavioral, EMG and EEG data from Experiment 1 (n = 13; mean ± SEM)

We used a slightly different baseline period for computing the burst threshold in our current study (prior to the Go signal; see *Extracting Beta Bursts* section) compared to our previous study (following the SSRT; Jana et al., 2019) in order to obtain more trials with bursts. This had a minimally different effect on the overall outcomes (see Table 1 and Jana et al. 2019). For the within-participant, trial-by-trial analysis we identified all Successful Stop trials containing both an EMG burst and a beta burst in the period before the EMG burst on that trial, and correlated the time of EMG cancellation (CancelTime) with the time of the beta burst (BurstTime) across all trials. Repeated measure correlation analysis showed that every participant had a positive slope in the relationship between BurstTime and CancelTime (Figure 2b). Moreover, this was significant at the group level in that there was a common positive relationship across individuals indicating that trials with later BurstTime were associated with later CancelTimes (r_m_ = 0.4; p< 0.001) (Figure 2b). This within-participant result is striking for scalp EEG which is typically a noisy metric. It highlights the value of studying beta bursts. We show that, within individual participants, at the single trial level, the timing of right frontal beta burst activity tracks the latency of stopping: later beta bursting is associated with slower stopping. These results provided guidance for our causal test of rIFC timing using TMS. Given that the mean BurstTime (reflecting the time of the peak amplitude of the beta burst) across participants was ∼125 ms (see Table 1), we estimated that the optimal time of the TMS pulse should be a little earlier than 125 ms such that it catches the rise of the burst. We therefore decided in Experiment 3 to stimulate with TMS at 80 ms after the stop signal.

**Figure 2.**
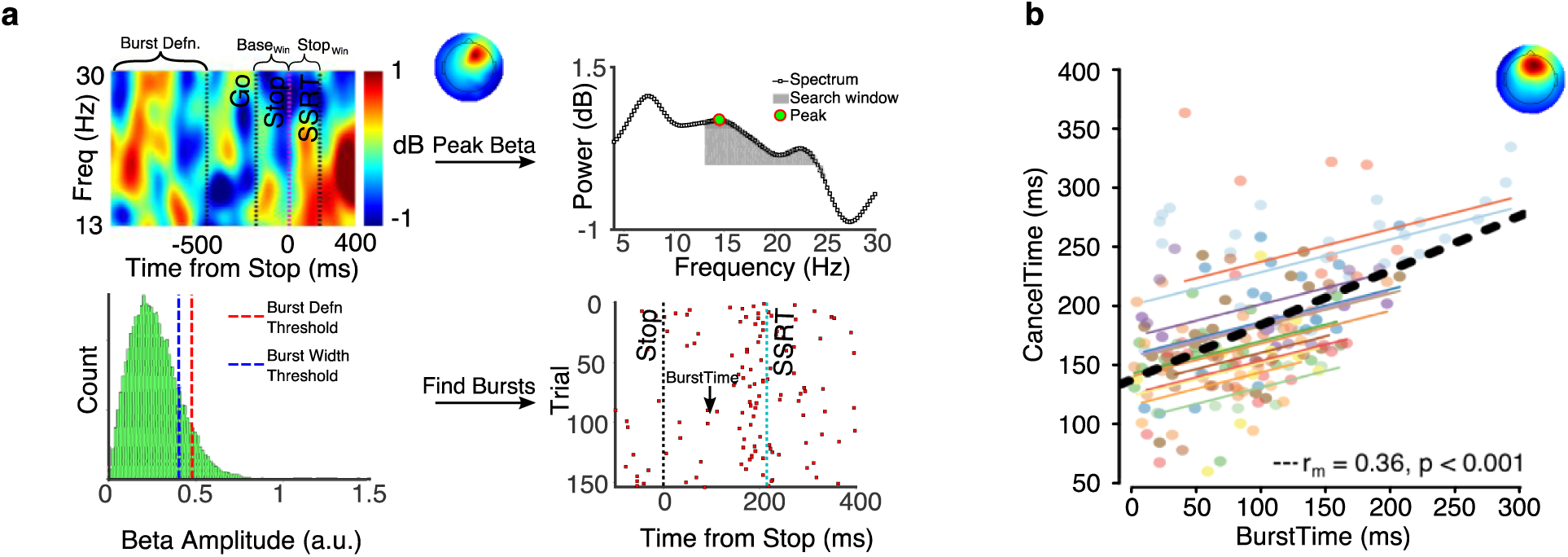
Experiment 1, beta bursts method and results. (a) Extracting beta bursts from a single participant. Top Left shows the average event-related spectral perturbation for the Successful Stop trials and the corresponding right frontal topography. The period between Go and Stop is the baseline window (Base_Win_) and from stop signal to SSRT is the stop window (Stop_Win_). The period before Go (−1000 ms to −500 ms) is the period taken to define the burst threshold. Top right is the peak beta power estimated in the Stop_Win_. Bottom Left is the peak beta power pooled across all trial types (Successful, Failed and Go) to define the burst threshold (dotted red line). Note that once a burst is detected using the burst definition threshold, we use the burst width threshold to compute the burst width and the burst probability across time. Bottom Right shows the raster plot for all trials for a participant, a red square denotes the peak of the burst which is defined as the BurstTime. (b) The trial-by-trial relationship between the BurstTime and the CancelTime within each participant. The colored lines show each participant’s relationship between BurstTime and CancelTime. The bottom right shows the common linear association obtained via the repeated measures correlation r_m_-value.

### Experiment 2 (fMRI and EEG experiment)

Our key aim for the major Experiment 3 below was to deliver temporally-precise TMS specifically to rIFC. As this required fMRI localization, we recruited a sub-sample (20) of 33 participants from a separate study wherein they had performed the Stop Signal Task on two separate occasions, once with fMRI and once with EEG. Here we re-analyze the EEG data to obtain an estimate of BurstTime in the same participants we would then run in Experiment 3 below. We start by briefly describing their stop signal behavior from the EEG session to highlight similar behavioral performance in the task across Experiments 2 and 3, some months apart. Next, we report the beta burst results. Please note that, unlike Experiment 1 above which had more than an hour of data per participant, the EEG stop signal data for Experiment 2 was only acquired in 12 minutes. Thus, we did not have enough trials to perform the within-participant single-trial BurstTime/CancelTime correlation analysis.

#### Stop Signal Task Behavior

Behavioral performance in the Stop Signal Task was as one would expect (Table 2), with Go response times being longer than Failed Stop response times, probability of stopping close to 50% and SSRT within 200-250 ms (Table 2). An EMG burst was identified in 67 ± 3 % of Successful Stop trials. The amplitude of these EMG bursts was ∼50% of that observed in Go trials, and CancelTime was ∼150 ms (Table 2).

**TABLE 2:**
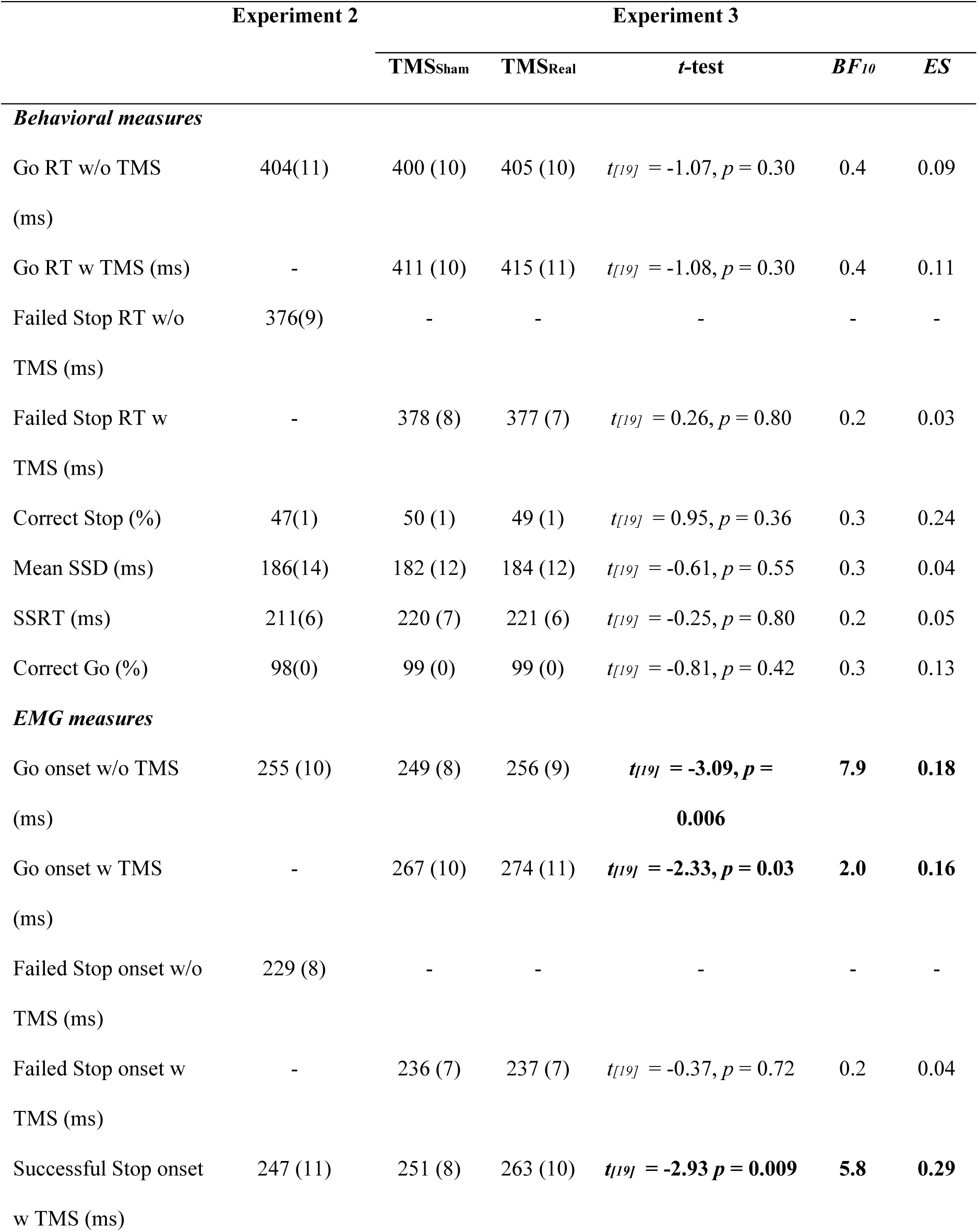
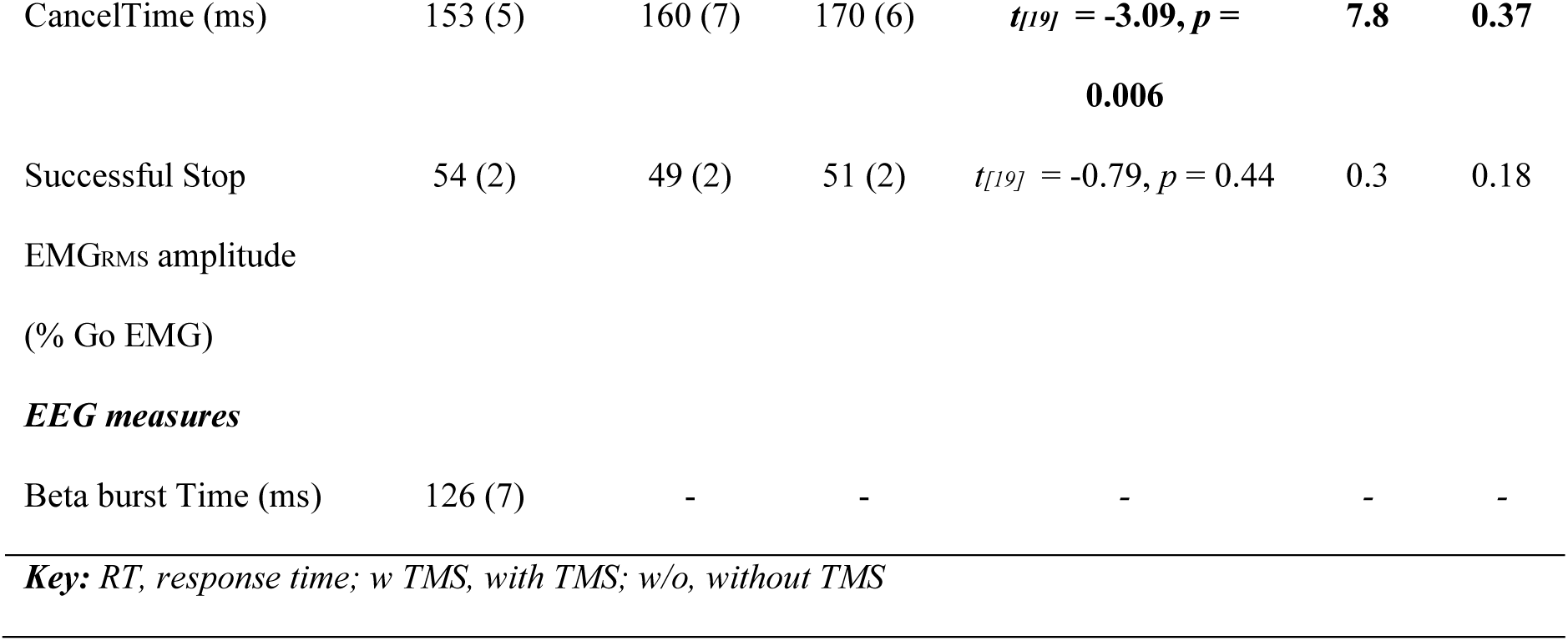
Behavioral, EMG and EEG data from Experiments 2 and 3 (mean ± SEM), along with test statistics comparing behavior for Real and Sham TMS in Study 3 (t-test, Bayes Factor and Effect Size). In Experiment 1 n = 20 and in Experiment 2 n = 20, 19 & 18 behavior, EMG & EEG.

#### EEG beta burst activity

We again analyzed the timing of right frontal beta bursts. In order to spatially localize these bursts, we first increased the signal to noise ratio for analysis by identifying a right frontal spatial filter for each participant, a proxy for the activity of the rIFC. For those participants who had a right frontal component (n = 18), we estimated the burst % in Successful Stop, Failed Stop and Correct Go trials. We saw a similar pattern of results for the burst % as for Experiment 1 (Figure 3a). The burst % for Successful Stop trials (22.5 ± 2.3 %) was greater than its baseline (13.0 ± 1.2 %; t_[17]_ = 5.0, p < 0.001, BF_10_ > 100) and the corresponding period in Go trials (15.5 ± 1.7 %; t_[17]_ = 3.8, p = 0.002, BF_10_ > 25.6). The average BurstTime (126 ± 7 ms) was also comparable with Experiment 1 and was significantly shorter than the mean CancelTime (156 ± 6 ms; t_[12]_ = 26.1, p < 0.001, BF_10_ > 100). Finally, we replicated the findings of Jana et al. (2019), showing that there was a significant positive correlation between the BurstTime and CancelTime across participants (r = 0.47, p = 0.048, BF_10_ = 1.5) (Figure 3b). These findings further validated our selected time of stimulation of the rIFC using TMS which we set as 80 ms. However, in this set of participants we were unable to interrogate the single trial relationship of BurstTime and CancelTime owing to the much lower number of trials as explained above.

**Figure 3.**
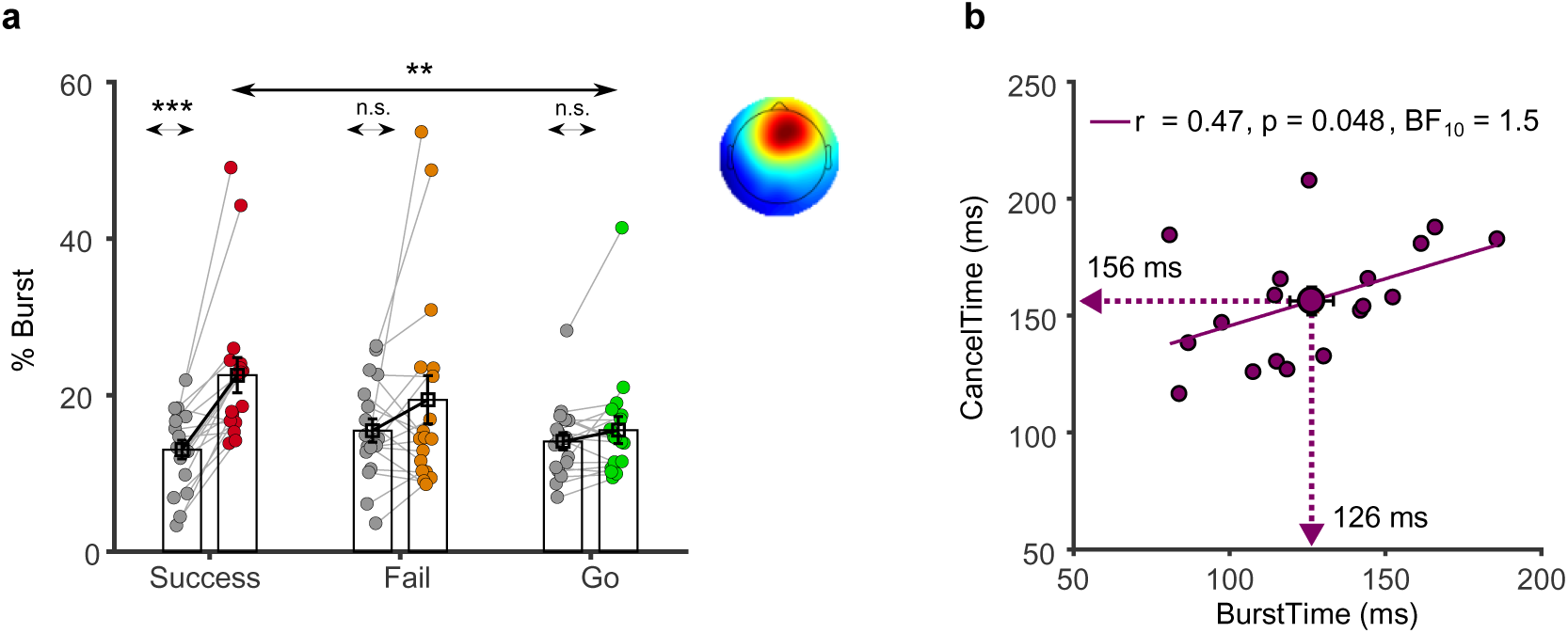
Experiment 2, beta burst results. (a) The burst % along with the average scalp topography across all participants in Experiment 2 for the Successful (pink), Failed (orange) and Go (green) trials in the Stop_Win_ compared to the Base_Win_ (grey). (b) The correlation between BurstTime and CancelTime across participants with the mean values represented by the arrow on each axis (* p<0.05, ** p< 0.01, *** p < 0.001).

### Experiment 3: TMS experiment

Having shown that the timing of right frontal cortex activity, indexed by the time of beta bursting, correlates with CancelTime at both the within- and between-participant level, we sought to examine a possible causal role of rIFC in stopping at this time. The same 20 participants from Experiment 2 returned for an experiment using TMS to provide a brief disruption to the rIFC at 80 ms after the stop signal, i.e. just prior to the average time of the beta bursting. Our dependent measure on each successful stop trial was CancelTime from the EMG.

Our pre-registered hypotheses were that, by comparison with TMS_Sham_, TMS_Real_ would: a) prolong measures of stopping latency, namely CancelTime and SSRT; b) result in increased EMG burst amplitudes on successful stop trials, since the bursts will have had longer to rise before being cancelled; c) not affect Go response times, assessed as both EMG onset and button presses. A final prediction was that the time at which TMS was given relative to implementation of the stop would influence the extent of stopping disruption.

The key finding, in keeping with our pre-registered hypothesis, was that TMS_Real_ was associated with a small increase in the latency of stopping compared to TMS_Sham_, albeit only for CancelTime and not for SSRT (Table 2, Figure 4a). TMS_Real_ did not affect the latency of response execution (Go or Failed Stop response times; Table 1, Figure 4a).

**Figure 4.**
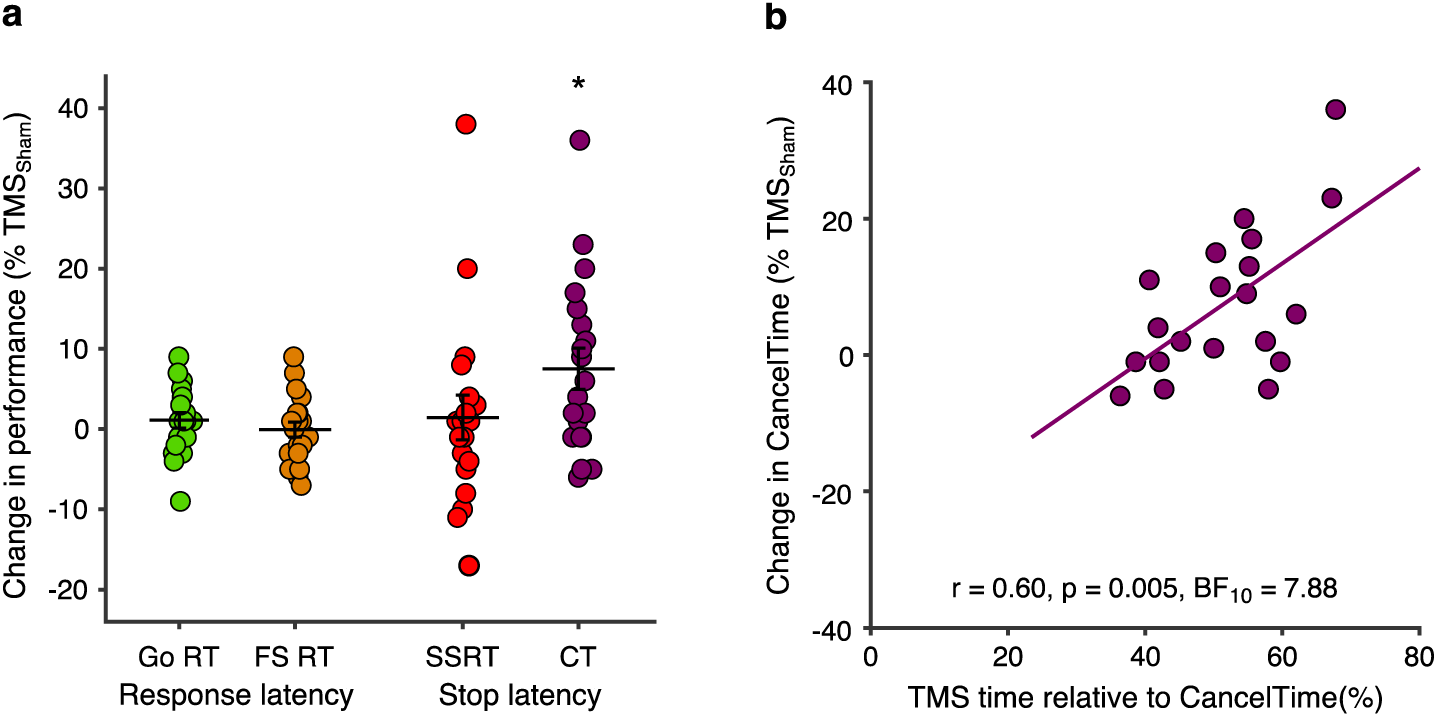
Experiment 3, TMS effects on behavior. (a) TMS_Real_ elongates CancelTime. Percentage change in response times (Go and Failed Stop) and stopping times (SSRT and CancelTime) for TMS_Real_ relative to TMS_Sham_, showing that only CancelTime was prolonged (*p < 0.05 one-sample t-test). (b) Participants who received TMS closer to the time of EMG decline showed the greatest increase in CancelTime. Pearson correlation between the percentage change in CancelTimes for TMS_Real_ relative to TMS_Sham_ and the relative time at which TMS was delivered with respect to CancelTime (where greater numbers indicate that TMS was delivered closer the end of CancelTime for that individual).

In the following sections we present a range of evidence supporting the idea that the prolongation of CancelTime specifically reflects an influence on the stop process by TMS_Real_ over rIFC, and is unlikely to be explained by other possible factors such as a non-specific effect of distraction or discomfort, or a slowing of EMG onset times.

#### Timing-specificity of the TMS effect on CancelTime

Our decision to deliver TMS 80 ms after the Stop Signal was motivated by the general agreement in beta burst timing across the different cohorts in Experiments 1 and 2 (∼125 ms after the Stop Signal). Although we did not have burst timings for Experiment 3, the general consistency of behavioral performance across sessions in Experiments 2 and 3 (typically <6 months apart) provides reassurance that the timing was appropriate (Table 1). Specifically, there were no statistically significant differences between Experiment 2 and the TMS_Sham_ condition in Experiment 3 for the major measures of Go and Failed Stop response times, mean stop signal delay, SSRT or CancelTime (all *t* ≤ 0.91, p ≥ 0.37, ES ≤ 0.18 and all BF_10_ ≤ 0.7).

Of course, the time of TMS was fixed across all participants and yet we know from previous work (Jana et al. 2019) and Experiment 2 that the timing of right frontal beta bursts varies across participants, and does so systematically with CancelTime. The implication is that the timing of TMS might not have coincided with right frontal beta bursting in all participants, and therefore might not have been optimal to disrupt stop-related processing in rIFC. This indeed seems to be the case, as we found a positive correlation between the percentage change in CancelTime for TMS_Real_ relative to TMS_Sham_ [calculated as: ((TMS_Real_ – TMS_Sham_)/ TMS_Sham_) × 100] and the relative time at which TMS was delivered with respect to the end of the Stop process (i.e. as a percentage of CancelTime in TMS_Sham_ trials; Figure 4b). In other words, those participants with a shorter CancelTime, and by implication an earlier beta burst time, received TMS closer to the end of CancelTime, and presumably closer to the time of beta bursting. These same participants showed the greatest prolongation of CancelTime in response to TMS_Real_. This is therefore an important post-hoc analysis that provides good evidence for the timing-specificity of the stopping disruption by TMS_Real_.

#### Further evidence that TMS disrupted stopping performance

While TMS_Real_ elongated CancelTimes it did not have an effect on SSRT, contrary to our pre-registered hypothesis (Figure 4a; Table 1). We therefore turned to a potentially more sensitive measure for evaluating the impact of TMS on behavior: examining Failed Stop response times as a function of the stop signal delay [see Bompas et al. (2019) for detail]. When a participant sees a stop signal, only the fastest responses escape inhibition (i.e. Failed Stop trials). Bompas et al. (2019), based on a distribution analysis over many trials, suggest that, for a particular stop signal delay, the time at which the Failed Stop diverges from Go trials is the earliest point at which stopping occurs. Here we looked at a subset of Failed Stop response times that: (i) had at least 5 observations for both TMS_Real_ and TMS_Sham_; and (ii) had a TMS pulse that occurred at least 60 ms before the response time (see Jana et al. for a discussion of why this corresponds to the key time period for rIFC mediated stopping). For each participant, we now calculated an aggregate measure across all stop signal delays of the median Failed Stop response time minus the stop signal delay, separately for TMS_Real_ and TMS_Sham_. We predicted that TMS_Real_ would elongate this measure. This behavioral analysis has the advantage over SSRT that it specifically looks only at the trials where the TMS pulse came early enough to have a causal effect on behavior (whereas the SSRT calculation uses all trials).

As expected, Failed Stop response times increased as a function of stop signal delay (Figure 5a). This illustrates that later stop signals result in a later divergence of the response time distribution, as increasingly later responses are allowed to escape inhibition via the putative stop process. In line with the TMS_Real_-related increase in CancelTime, stopping latencies estimated as the mean Failed Stop response time minus the mean stop signal delay tended to be greater for TMS_Real_ (202 ± 19 ms) compared to TMS_Sham_ (195 ± 21 ms), but the difference did not cross the threshold for statistical significance (paired *t*-test: *t* = 1.89, *p* = 0.073, *BF_10_* = 1.03). A one sample *t*-test on the percentage change for TMS_Real_ relative to TMS_Sham_ was also close to the threshold for statistical significance (mean change: 4 ± 8%; one-sample *t*-test: *t* = 2.02, *p* = 0.057, *BF_10_* = 1.25). We now tested whether, across participants, there would be a correlation between this Failed-Stop derived measure of inhibition latency and CancelTime. There was indeed a significant positive correlation (Figure 5b). These results therefore buttress our finding that TMS_Real_ elongated CancelTime by also showing an effect on behavior (when looking specifically at the trials where the TMS pulse came early enough, and using an aggregate measure of the earliest inhibition latencies for participants).

**Figure 5.**
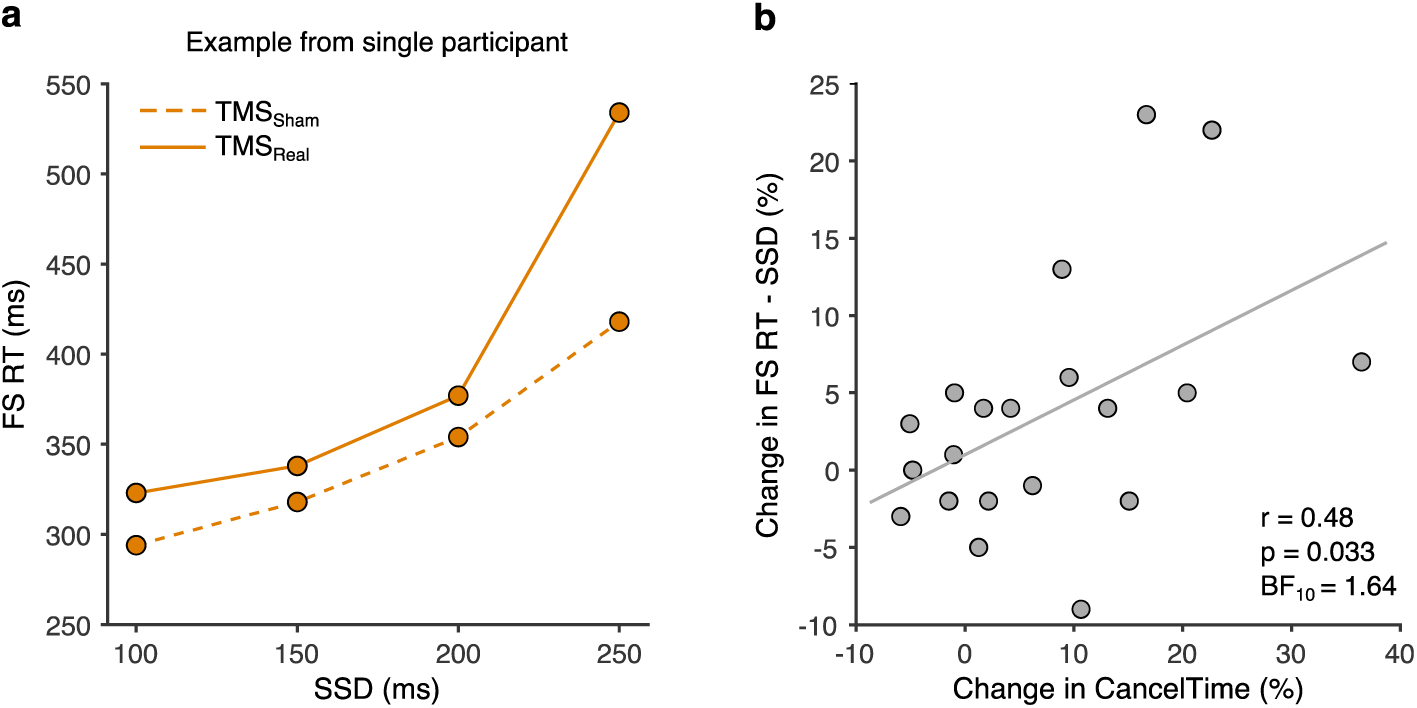
Experiment 3, TMS effects on Failed Stop response times as a proxy measure of stopping latency. (a) Longer stop signal delays result in later intersection of the response time distribution by a presumed inhibitory process. Representative data from a single participant illustrating that Failed Stop response times increase as a function of stop signal delay. Note too, that Failed Stop response times were longer for TMS_Real_ compared to TMS_Sham_. (b) Participants showing a greater increase in CancelTime for TMS_Real_ compared to TMS_Sham_ also showed a greater increase in the Failed Stop-derived measure of stopping latency, calculated as the mean Failed Stop response time minus the mean stop signal delay.

#### Efficacy of sham control and double blinding

TMS over inferior frontal areas of the scalp is well known to be uncomfortable/distracting (Meteyard and Holmes 2018), because of more facial muscle twitching and possibly auditory input produced by the TMS click. Moreover, the extent of this discomfort/distraction is related to the degree of behavioral impairment in cognitive and motor tasks (Abler et al. 2005; Meteyard and Holmes 2018). As described in the methods, we used a realistic sham coil (which produces both sound and facial twitches) as a comparison for real TMS in order to control for such non-specific effects of stimulation. What is key is that, within the double-blind methodology, we calibrated the TMS_Sham_ stimuli at the start of the experiment to achieve a similar perceived intensity of stimulation as TMS_Real_. The efficacy of the TMS_Sham_ in mimicking the sensations of TMS_Real_, and their associated effects on behavior, is supported by the fact that the prolongation of Go response times for TMS trials compared to No-TMS trials was similar for TMS_Real_ and TMS_Sham_ (Table 1; rmANOVA: TMS coil, F_[1,19]_ = 1.27, p = 0.27, *η_p2_* = 0.06; TMS presence F_[1,19]_ = 36.17, p < 0.001, *η_p2_* = 0.66; TMS presence × TMS coil, F_[1,19]_ = 0.03, p = 0.86, *η_p2_* = 0.002).

Although we attempted to match the perceived intensity of TMS_Real_ and TMS_Sham_, 13/20 participants still reported noticing changes in the sensations throughout the experiment, and cited differences in the perceived intensity (13/13) and location (4/13; e.g. more eye or facial muscle twitching). Nevertheless, there was no clear difference in the prolongation of CancelTime between participants who did report noticing changes in the stimulation during the experiment (11.4 ± 5.4 % change in CancelTime) and those who did not (5.4 ± 2.3 %; two-sample t-test, *t_[18]_* = −1.20, *p* = 0.24, ES = 0.51 and BF_10_ = 0.67).

The experimenter correctly identified the coil being used (TMS_Real_ or TMS_Sham_) in a given set of blocks in 8 out of 20 experimental sessions (40%). This is below the level of chance, indicating that the experimenter was effectively blind to the TMS coil being used. In addition, there was high consistency in the positioning and orientation of the TMS coil (i.e. small errors with respect to the target) throughout the experiment: target, 0.7 ± 0.03 mm; angular, 6 ± 0.2°; and twist, −0.8 ± 0.1°.

#### Prolongation of CancelTime is unrelated to delayed EMG onset

A potential point of concern when interpreting the increase in CancelTime for TMS_Real_ is that it could simply be a by-product of the slowed EMG onset times observed for TMS_Real_ compared to TMS_Sham_ in both Go and Successful Stop trials (Table 2). In other words, it might simply indicate that EMG bursts were delayed on the whole, with both their onsets and declines shifted later in time. However, we do not think that the shift in EMG onset and decline times are related. Firstly, the timing of the EMG decline coincides quite tightly with a broad motor system suppression (Jana et al. 2019), implying that it results from an active withdrawal or suppression of motor drive and is thus not time-locked in any way to the onset of EMG. Secondly, the elongating effect of TMS_Real_ on CancelTimes displayed a degree of timing specificity (see above; Figure 4b), whereas percentage change in Successful Stop EMG onset times was unrelated to the relative timing of TMS (*r* = −0.05, *p* = 0.84, *BF_10_* = 0.17). Third, there was no relationship between the percentage change in CancelTimes and the percentage change in Successful Stop EMG onset times across coils (*r* = 0.26, *p* = 0.26, *BF_10_* = 0.32). Fourth, participants with an above-median percentage change in Successful Stop EMG onset times had CancelTimes that were statistically similar to participants with a below median change (7.5 ± 2.5 % versus 7.5 ± 4.4 % change in CancelTime; two-sample *t*-test: t_[18]_ = 0, p = 1.0, BF_10_ = 0.31).

## DISCUSSION

The current experiments tested the idea that rIFC plays a critical role in human action-stopping, and more specifically that the efficacy of stopping is sensitive to the timing of rIFC activation. We focused particularly on Successful Stop trials with small bursts of EMG activity from which we derive our pseudo-single trial estimate of stopping latency, CancelTime. Experiment 1 showed that CancelTime correlated to the timing of bursts of right frontal activity in the beta band (BurstTime) at the single trial level: trials with earlier beta bursts were associated with earlier stopping. This result is highly consistent with the idea that the stop process is initiated via the right prefrontal cortex. This result is also striking for being a rare demonstration of within-participant, single-trial, EEG correlation with executive function. A separate cohort participated in experiments 2 (MRI and EEG) and 3 (TMS). From the Experiment 2 data we replicated the finding that the timing of right prefrontal beta bursts following a stop signal is ∼125 ms, and thus 40-50 ms prior to CancelTime. This motivated the timing of our disruptive TMS pulses in Experiment 3. TMS pulses delivered over rIFC 80 ms after the stop signal, and thus just prior to BurstTime, prolonged CancelTime. Furthermore, this disruption was most pronounced in participants for whom the pulse appeared closer to the time of EMG cancellation. Overall, our results help refine the timing of a putative prefrontal-basal ganglia-thalamocortical network for action-stopping (Aron et al. 2014; Jana et al. 2019), wherein rIFC plays a causal role in the early stage of implementing the stop process.

Our key result was that TMS delivered to right pars opercularis ∼45 ms prior to the mean BurstTime had the effect of increasing CancelTime, but left response execution times in Go and Failed Stop trials unaffected. This specific effect on stopping implies that the TMS pulse briefly disrupted processing in rIFC that was functionally relevant to cancelling the impending action. Moreover, we also saw some evidence for a timing-specific, causal role of rIFC. Specifically, we showed that the people with a shorter CancelTime evidenced the greatest TMS-induced disruption in CancelTime (Figure 4b). We speculate that right prefrontal beta bursts, TMS time and CancelTime are related in the following way. First, we know from previous work that participants with a longer BurstTime also have a greater CancelTime (Jana et al. 2019). It follows that participants with a shorter CancelTime would have received TMS closer to BurstTime as TMS was always delivered at 80ms after the stop signal. Since the effect of TMS on behavior is often short-lived (e.g. tens of milliseconds; Amassian et al., 1989; Pitcher et al., 2008), it might just have lasted long enough to disrupt stop-related right prefrontal cortex activity in those with shorter CancelTimes, but would have occurred too soon to affect it in those with longer CancelTimes. Indeed, one might speculate that had we adjusted the time of the TMS pulse according to each individual’s BurstTime or CancelTime that we might have seen a bigger disruption in stopping latency at the group level.

These experiments have important implications for the study of basic action-stopping, and more broadly for understanding individual differences in impulsivity. Firstly, our pseudo-single trial measure of stopping latency, CancelTime, was fundamental to establishing the trial-by-trial relationship between beta and stopping. This would not have been possible with the standard measure, the SSRT, which reflects a single (over-)estimate (Jana et al. 2019; Skippen et al. 2019) of stopping latency per person. Secondly, we have shown the value of using beta bursts to reveal the fine timing of stopping related brain activity (see also Wessel, 2019). Taken together, these methods provide a good basis for testing whether the variation in sub-processes of action-stopping might better relate to real-world self-control and impulsivity than SSRT (Friedman and Miyake 2004; Lijffijt et al. 2004; McLaughlin et al. 2016; Chowdhury et al. 2017; Enkavi et al. 2019; Skippen et al. 2019).

Nonetheless there were a few puzzles. One is that we originally predicted that SSRT would also be prolonged by TMS and given that CancelTime and SSRT are strongly correlated (Jana et al. 2019), it is puzzling that it was not. One explanation is that CancelTime reflects a more direct measure of stopping latency that is less sensitive to the variability of response times, which encompass large electromechanical delays (∼100 ms between EMG onset and button press; Jana et al., 2019), that otherwise influence the calculation of SSRT. Indeed, when we used a potentially more sensitive measure of behavioral stopping latency we did see an effect of TMS_Real_. Specifically, when looking at stop trials where the TMS came early enough to potentially influence stopping, and using the aggregate measure of Failed Stop response time minus stop signal delay (as a proxy of the earliest inhibition latencies), we found a marginally significant elongation of this measure for real versus sham. More compellingly, across participants this delta was correlated with the delta in CancelTimes.

Another open question concerns the spatial specificity of the effect of TMS on stopping. Ideally one would add another condition with real and sham TMS on different brain site. However, there simply wasn’t enough time in a session to do all these conditions together. Note that there were only 25% stop trials of which 50% were successful and of these, only ∼50% contained EMG bursts; and this had to be repeated for TMS_Real_ and TMS_Sham_. Consequently, we do not know whether similar results would have been achieved stimulating other prefrontal cortical areas such as the pre-supplementary motor area, which is considered part of the stopping network (Cai et al. 2012; Coxon et al. 2016; Jahanshahi and Rothwell 2017; Kohl et al. 2019), or the inferior frontal junction, which is thought to regulate attentional capture during stopping (Verbruggen et al. 2010; Aron et al. 2014; Sebastian et al. 2016). Indeed, it is even possible that the disruption in rIFC spread to functionally-connected areas such as the pre-supplementary motor area, and that this contributed to changes in stopping latency. If this were true, the current results would still indicate that rIFC, along with the wider stopping network, is engaged in implementing the stop at this time. Finally, it remains open whether the right prefrontal beta bursts are causally relevant to stopping. Specifically, we do not know whether the prolongation of CancelTime was because we disrupted ongoing beta, or whether we disturbed some other process coinciding temporally with the beta bursts. Here we assumed only that frontal beta bursts provided a marker of when prefrontal cortex was actively involved in processing relevant to stopping. Future studies might address this issue by causally modulating frontal beta, perhaps endogenously via neurofeedback (Khanna and Carmena 2017) or exogenously via brain stimulation (Hanslmayr et al. 2014), and evaluating its subsequent effect on the suppression of actions.

In conclusion, we show, first, that action-stopping is preceded by beta bursts over putative rIFC, and that the timing of these bursts correlates with the latency of stopping at a single-trial level so that earlier bursts were associated with faster stopping. Next, we showed that TMS over rIFC prolonged stopping latencies and, moreover, the prolongation was most pronounced in individuals for whom the pulse appeared closer to the presumed time of beta bursting. These results help validate a prominent model of the neural architecture of action-stopping, whereby the process is initiated early by the rIFC (∼80-120 ms after a stop signal) and is then implemented via basal ganglia and primary motor cortex, before affecting the muscle at about 160 ms. The results also highlight the usefulness of prefrontal beta bursts to index an apparently important sub-process of stopping, the timing of which might help explain within- and between-individual variation in impulse control.

## NOTES

URL #1 available at: osf.io/93rb7

URL #2 available at: osf.io/b2ng5

## Acknowledgements

We gratefully acknowledge funding support from NIH NS106822, DA026452 and the James S McDonnell Foundation 220020375.

